# Innate heuristics and fast learning support escape route selection in mice

**DOI:** 10.1101/2021.12.14.472555

**Authors:** Federico Claudi, Dario Campagner, Tiago Branco

## Abstract

When faced with imminent danger, animals must rapidly take defensive actions to reach safety. Mice can react to innately threatening stimuli in less than 250 milliseconds [1] and, in simple environments, use spatial memory to quickly escape to shelter [2,3]. Natural habitats, however, often offer multiple routes to safety which animals must rapidly identify and choose from to maximize the chances of survival [4]. This is challenging because while rodents can learn to navigate complex mazes to obtain rewards [5,6], learning the value of different routes through trial-and-error during escape from threat would likely be deadly. Here we have investigated how mice learn to choose between different escape routes to shelter. By using environments with paths to shelter of varying length and geometry we find that mice prefer options that minimize both path distance and path angle relative to the shelter. This choice strategy is already present during the first threat encounter and after only ~10 minutes of exploration in a novel environment, indicating that route selection does not require experience of escaping. Instead, an innate heuristic is used to assign threat survival value to alternative paths after rapidly learning the spatial environment. This route selection process is flexible and allows quick adaptation to arenas with dynamic geometries. Computational modelling of different classes of reinforcement learning agents shows that the observed behavior can be replicated by model-based agents acting in an environment where the shelter location is rewarding during exploration. These results show that mice combine fast spatial learning with innate heuristics to choose escape routes with the highest survival value. They further suggest that integrating priors acquired through evolution with knowledge learned from experience supports adaptation to changing environments while minimizing the need for trial-and-error when the errors are very costly.

## Results

### Escape route choice is determined by path distance and angle to shelter

To investigate escape route choice, we placed mice in elevated arenas with a threat and a shelter platform connected by runways of different configurations and lengths. Previous work has shown that in simple arenas mice escape along a direct vector toward a memorized shelter location [2,7,8], and that they can form subgoal memories to avoid obstacles when navigating to shelter [3]. To determine whether mice learn the value of alternative routes to shelter, we first built an arena where the direct path to shelter lead to a dead-end and while two other open paths of equal length were available (Figure 1A). Mice explored the entire arena over a period of ~10 minutes (Figure S1A), after which they were exposed to innately threatening auditory and visual stimuli [8,9] when they were on the threat platform, facing away from the shelter. Mice reliably escaped from threat in these conditions (Figure S1B) and had a preference for the two side paths (P_side path_ = 0.79, P_dead-end_ = 0.21, 34 trials from 7 mice; Figure 1A; Video 1). The average time to leave the threat platform to one of the side arms was 2.49±0.81s, and mice accelerated directly toward the left of right paths from the start of the flight (Figure 1B, 1C). This shows that mice quickly learn to overcome the innate preference of escaping along the shelter direction and use knowledge of which paths lead to shelter when committing to escape trajectories.

**Figure 1.**
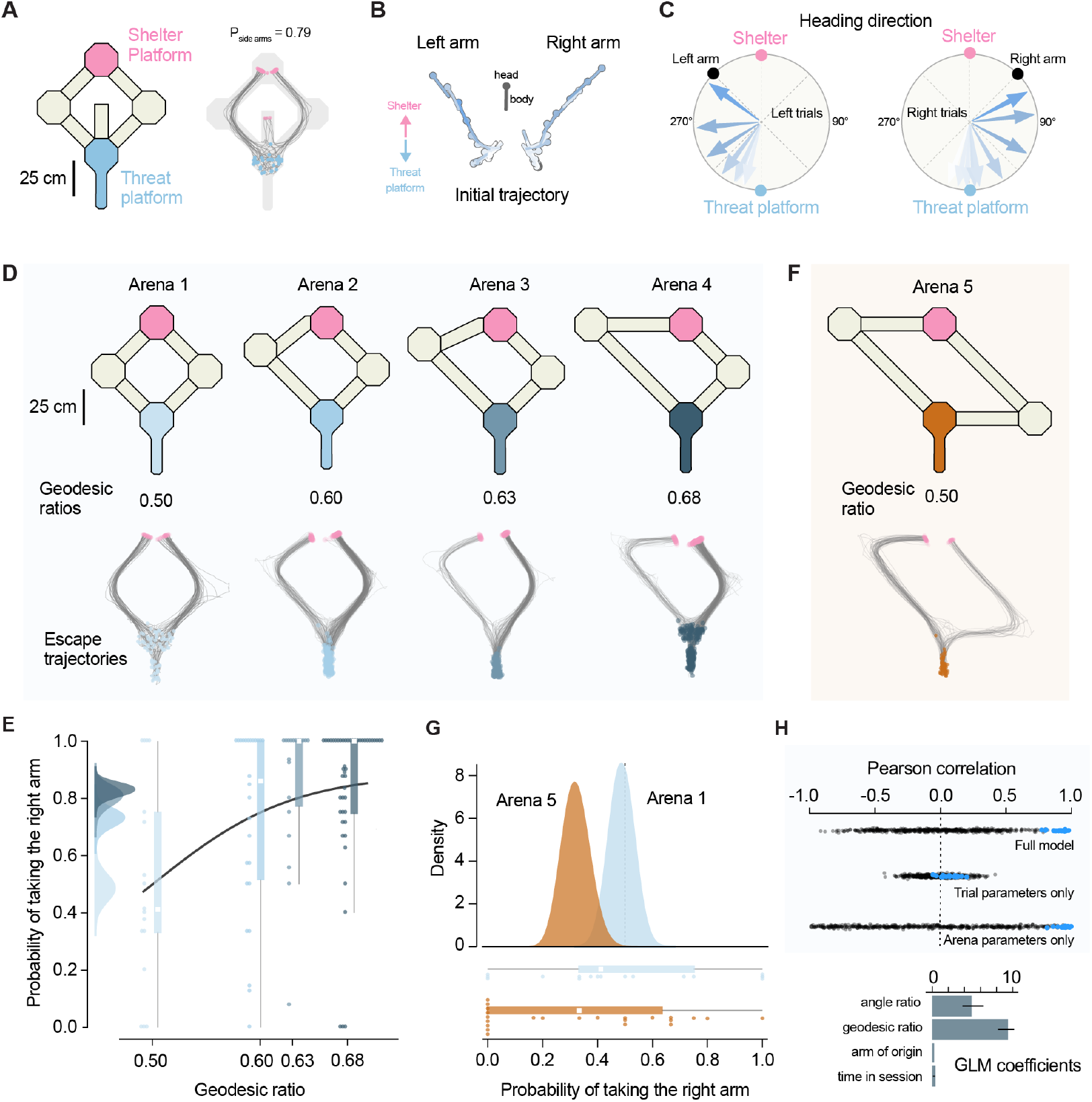
Escape route choice is determined by path distance and angle to shelter. **(A)** Left, schematic view of Arena 1 with a dead-end central arm. Right, movement tracking traces for all escape trials (grey). Blue and pink circles mark the start and end of each escape run, respectively (p=0.038, binomial test for P_dead-end_ = 0.33). **(B)** Example trajectories on the threat platform for two escapes initiated in the left and right arm. The position of the head and body of the mouse is shown at 250 ms intervals. Color shows time elapsed from stimulus onset (later time points have darker shades of blue). **(C)** Average heading direction on the threat platform for left and right escapes. Each arrow shows the average heading directions at eight time points equally spaced between stimulus onset and exiting the threat platform (pooled across trials and animals). Color shows time elapsed from stimulus onset (later time points have darker shades of blue). **(D)** Top, arena schematic and corresponding geodesic ratio. Bottom, tracking traces from all trials in each arena with starting (blue) and end (pink) locations shown. **(E)** Probability of escape along the right path (P_right_.) in arenas 1 to 4. Scatter dots are P_right_ of individual mice and box plot shows median and interquartile range for all trials pooled for each arena. The left panel shows the posterior distributions of P_right_ from the Bayesian model (see Methods). **(F)** Top, schematic view of Arena 5, with the same geodesic ratio of arena 1 but with a different angle ratio between the two arms. Bottom, tracking traces from all trials in Arena 5 with start (orange) and end (pink) locations indicated. **(G)** Top, posterior distribution for P_right_ computed with the Bayesian model for all trials in Arenas 1 and 5. Bottom, P_right_ of individual mice (scatter dots) with median and interquartile range for pooled data. **(H)** Top, cross-validated Pearson correlation between predicted P_right_ and observed choice behavior. Data shown for the full model and two partial models - trial parameters only (arm of origin and time) and arena parameters only (arm length and angle). Blue dots are fits to the data, black dots are fits to shuffled data. Bottom, coefficient weights for the four predictor variables included in the GLM (mean and standard deviation over repeated tests, see Methods).

Next, we tested escape route selection in four different arenas where the length of the left arm was progressively increased while keeping constant the initial angle between each path and the threat platform (arenas 1-4, Figure 1D). In this experiment the relative value of the left hand-side path decreases between arenas 1 and 4 as mice should in principle escape along the shortest path to minimize exposure to danger [10–12]. The differences in geodesic distance (path length) between the threat and shelter platforms translated not only into differences in distance travelled but also into time taken to traverse each path during escape (Figure S1C, D). While path choice was probabilistic, when presented with threat, mice preferentially escaped via the shortest path in each of the three asymmetric arenas (overall P_right path_ = 0.79, 600 trials from 83 mice; Figure 1D,E; Video 2). Across all arenas, the probability of taking the shortest path was significantly dependent on the geodesic distance ratio between the two paths (arena1 P_right path_ = 0.486, arena 2 P_right path_ = 0.731, arena 3 P_right path_ = 0.807, arena 4 P_right path_ = 0.829, p = 0.0006 *one-way ANOVA*). The preference for the shortest path could not be explained by differences in arm familiarity arising from biases during arena exploration, nor by mice simply choosing the arm they entered the threat platform from (Figure S1E,F). This suggests that mice are choosing to escape along the shortest path. In addition, the time to leave the threat platform was independent of path chosen (2.53±1.51 and 2.58±1.37s for left and right arm respectively) and path choice could be predicted from the escape trajectory before mice left the threat platform (Figure S2). This indicates that mice commit to one path from escape onset. These data therefore suggest that mice evaluate either geodesic distance or escape duration to shelter when choosing escape routes. These two quantities are strongly correlated in our experimental setup (Figure S1C) and we cannot disambiguate between these two alternatives. In either case, however, mice quickly learn to select the fastest escape routes to safety.

To assess whether other aspects of the path geometry influenced escape route selection, we next built an arena where the two arms had the same length but the initial angle relative to the shelter was larger for the right arm (Figure 1F). In this configuration, when mice escape along the right arm, they initially must navigate away from the shelter direction, but the path length and escape duration is the same for both arms (Figure S1C). Thus, if mice selected escape paths based on path length or travel duration alone, they should choose each path with equal probability, as in arena 1. Instead, we found that mice had a clear preference for the left arm (P_left path_ = 0.69, 81 trials, 23 mice, p=0.02 *chi squared test*, Figure 1G). This suggests that in addition to geodesic distance, mice also consider shelter direction when choosing escape paths. To further quantify the relative weights of different variables on escape path preference we used a generalized linear model (GLM) to predict escape path choice across trials. The predictors included geodesic distance and shelter angle, as well as exploration time and trajectory before threat. The model fitted to the data could explain more than 90% of the variance in path choice in cross-validated tests and showed approximately equal weighting between geodesic distance and shelter angle, with minimal weight for exploration time and arm of origin (Figure 1H). Together these results show that when faced with two possible paths to shelter, mice can quickly learn distances and angles to shelter and escape along the route that minimizes both.

### Route learning does not require escape experience and is flexible

Computing escape route choices requires at least two steps: learning the properties of the available paths to safety and a function to map those properties into their value for escape (e.g.: favoring escape along the shortest path). While the first is likely learned during natural exploration of the environment, the second could in principle be learned through repeated encounters with threat, or it could be innate (i.e.: the animal is born with a value function that links path properties to escape values). To distinguish between these two alternatives, we computed path choice probabilities for the first trial of threat presentation (naïve) and compared them with the probabilities for trials after experience with threats (experienced). We found that the preference for shorter paths and smaller angles to shelter was already present in naïve trials, and that the path choice probabilities were not different between naïve and experience trials (Figure 2A). In addition, choice probabilities did not significantly change over the course of the experimental session and repeated threat presentations (Figure 2B), which is also in agreement with the low GLM weighting for the time in session variable (Figure 1H). This analysis suggests that the strategy for selecting escape routes does not develop through experience of escaping. Instead, the evaluation of path length and angle to shelter represents an innate heuristic to assign escape value to the different route options. In addition, the preference for selecting the shortest path upon the first exposure to threat implies that mice were able to learn the relevant environment properties during natural exploration of the arena. In our experiments, mice spend on average 11.03±3.8 min exploring before threat presentation, during which time they perform only 4.2±0.9 complete trips between the threat platform and the shelter (Figure 2C, D). Mice thus require a very small amount exploration to learn spatial properties relevant for escape and have an innate function to assign escape value to alternative paths based on the learned spatial relationships between the paths and shelter location.

**Figure 2.**
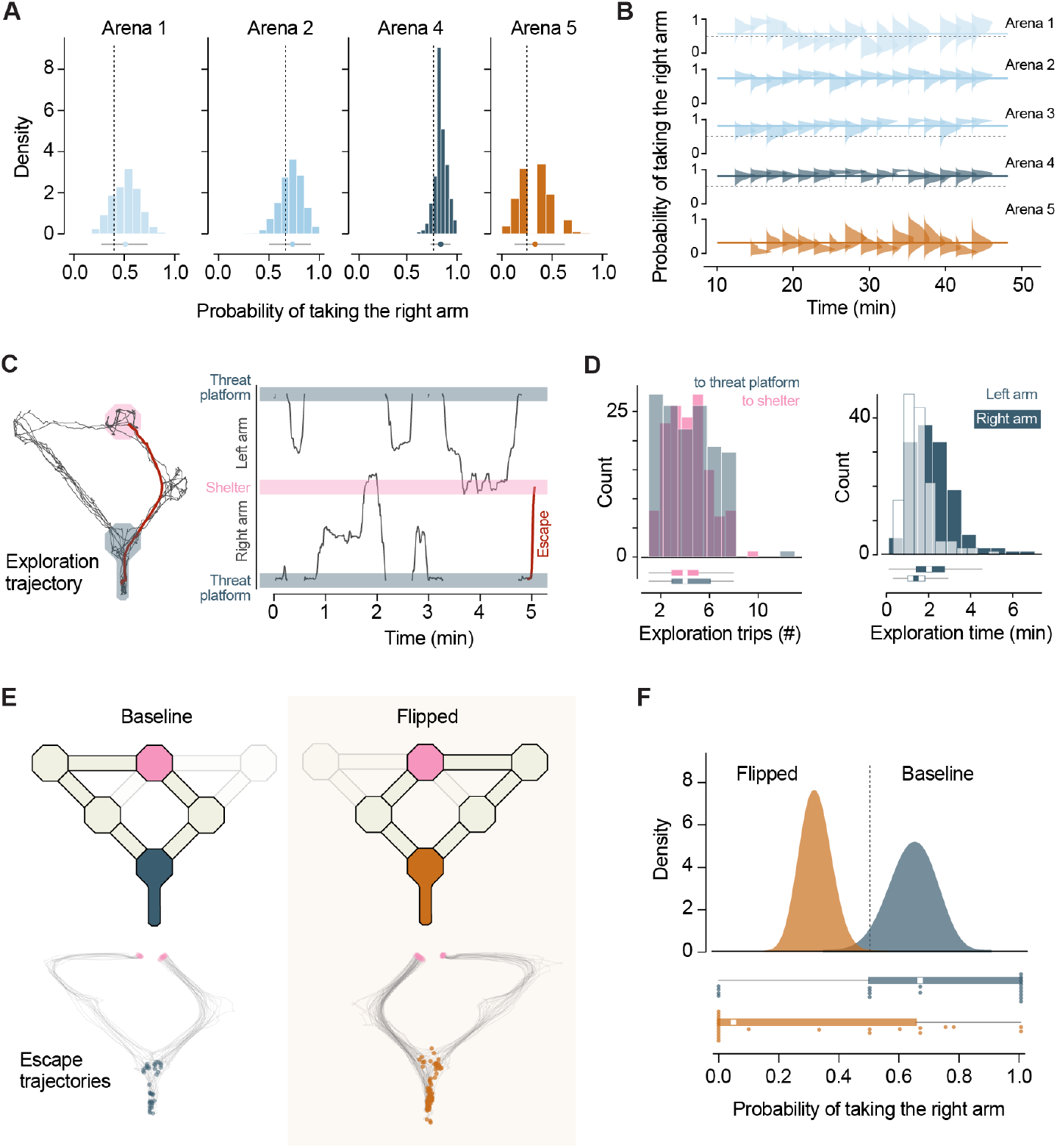
Route learning does not require escape experience and is flexible. **(A)** Distributions of P_right_ for data subsets randomly sampled from all experienced trials in each arena (mean and 95th percentile confidence shown underneath). Dashed line shows P_right_ for naïve pooled across mice. **(B)** Change in P_right_ over time (within single experimental sessions). The posterior distribution of P_right_ calculated from trials binned by time in the experiment is shown for arenas 1-5 (trials pooled across animals). Solid line shows the P_right_ for the entire duration of the session. **(C)** Left, example movement trajectory during arena exploration. Right, the same trajectory on the left, linearized to show the position of the mouse in the arena overtime. Red trace shows the trajectory for the first escape following the exploration period. **(D)** Left, histogram for the number of shelter-to-threat and threat-to-shelter trips during exploration across all experiments in Arenas 1-5. Right, histogram for total time exploring the left and right arms, pooled across all arenas. **(E)** Top, schematics of the dynamic arena in baseline and flipped configurations. Bottom, movement tracking trajectories for escapes in the baseline and flipped conditions (blue and orange dots show initial location, pink dots show final position). **(F)** Top, Bayesian model posterior estimates of P_right_ for trials from the baseline and flipped conditions. Bottom, scatter dots show P_right_ for individual mice, box plot shows median and interquartile range for pooled trials.

The combination of rapidly learning path properties and having an innate value function allows mice to effectively select escape paths shortly after entering a novel environment. Next, we aimed to establish whether this can also support flexible and adaptive escape route selection in a dynamic environment, where path preference must be rapidly updated to reflect changes in the arena. We built a version of arena 4 where the path lengths could be quickly and automatically flipped between left and right sides (Figure 2E; Video 3). After exploration and 2-3 threat presentation trials, we flipped the arena and let the animals again explore the maze (14.6±6.1 min, median, 6.05±2.77 threat platform to shelter trips). We then presented threats and found that the path preference during escape changed to reflect the new arena geometry – mice now took the left arm with a higher probability (baseline P_right path_ = 0.641, flipped P_right path_ = 0.321, p=0.0014 *Fisher’s exact test;* Figure 2F), while the time out of threat platform and orientation movement profiles were similar between baseline and flipped trials (2.16±1.10 and 1.77±0.93 seconds respectively, p=0.06 *t-test*). These data suggest that after initially learning the arena geometry and developing an escape route preference, mice remain in a flexible learning state where they can incorporate new information at a rate similar to naïve animals. This ability enables the selection of the fastest escape routes in changing environments.

### Model-based reinforcement agents with limited experience choose the shortest escape route

Learning the shortest escape route in our experiments was a fast process, which contrasts with large the amount of training needed for some spatial navigation and decision making tasks [13,14], as well as for training artificial intelligence agents [15,16]. To gain further insight into the type of learning algorithms that mice might be using, we compared the performance of different reinforcement learning (RL) algorithms [17] on a task similar to our experiments. We selected three algorithms representing different classes of RL models: Q-learning (model-free; [17]), DYNA Q (model-based; [17]) and influence zones (IZ; [18]). The latter is a model-based algorithm where several state-action values are updated simultaneously according to a topological mapping between states, and thus particularly appropriate for spatial navigation tasks [18]. These models were trained to navigate a grid-world representation of arena 4, from a starting location (corresponding to the threat platform in our experiments) to a goal where they received a positive reward (corresponding to the shelter location, Figure 3A). As the goal of the agents is to maximize the time discounted cumulative expected reward, this should result in learning a policy that selects the shortest route to the goal, thereby mimicking innate preference of mice for selecting shorter escape paths.

**Figure 3.**
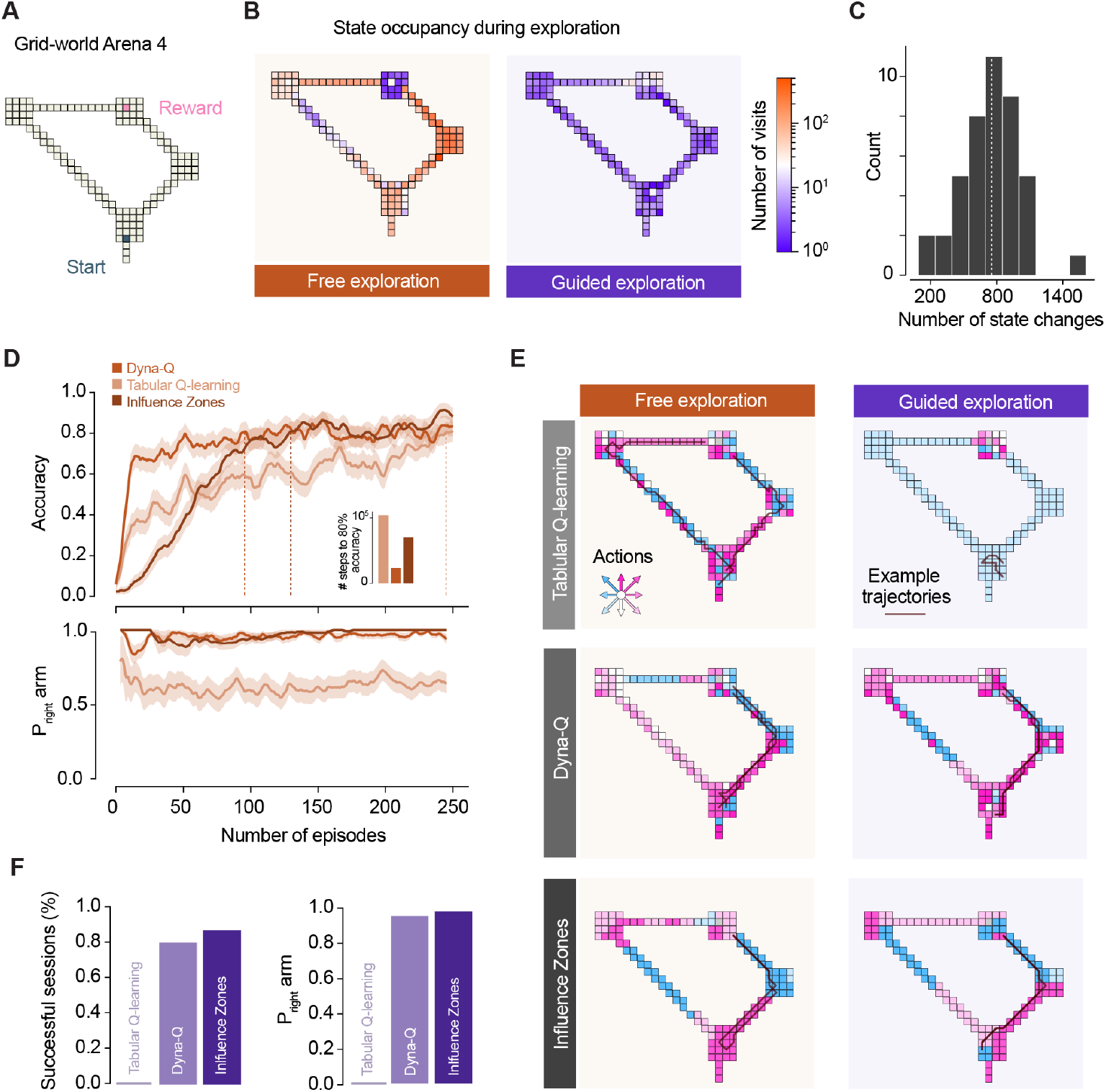
Model-based reinforcement agents with limited experience choose the shortest escape route. **(A)** Schematic of the grid world arena used for RL simulations. **(B)** Heatmap for the number of visits to each state during ε-greedy (left) and guided (right) exploration (data from two representative simulations). **(C)** Distribution of number of state changes during guided exploration across sessions. **(D)** Learning curves for simulations under ε-greedy exploration. Top, accuracy for the different model classes tested (fraction of agents that reaches the goal state during the evaluation trial; traces are mean and standard error of the mean across multiple model instances. Dotted lines mark when 80% success rate is reached, inset shows number of training steps to reach 80% accuracy. Bottom, probability of choosing the right arm in successful trials during training, for each RL model class. **(E)** Illustration of the policies for the different RL simulations after training. Inset arrows show all possible actions and the respective colors are shown in the arena to represent the best action that each class of RL models learned for every state in the arena. Lines show two example trajectories from trained agents attempting to navigate from the start to the goal location. **(F)** Left, performance of agents trained under the guided exploration regime. Left, outcome (success or failure) for each class of RL algorithm across 42 sessions. Right, probability of taking the right arm in successful sessions.

We trained the RL agents under two regimes: in the *free-exploration* regime, agents were allowed to explore the environment freely under an epsilon-greedy policy for 250 episodes with a maximum of 500 steps each; in the *guided-exploration* regime, RL agents explored the environment in a single episode and moved through the maze following the exploration trajectory of individual mice recorded during our experiments (see methods). The *free-exploration* regime is therefore analogous to the standard practice in the RL field [17] where agents are allowed a large number of steps to learn (up to 125,000 in our conditions). The *guided-exploration* regime poses a more challenging learning problem in principle: real exploration trajectories in our experiments have a mean of 754±267 steps, almost three order of magnitude less (Figure 3B, C). Under *free-exploration*, all models successfully learned to navigate to the goal location (Figure 3D). The short arm was chosen by 64% of the agents trained with Q-learning that reached the shelter at the end of the test trial, and by more than 95% when trained with DYNAQ or influence zones. (Figure 3E). In contrast, under *guided-exploration*, the Q-learning algorithm failed to learn how to reach the goal entirely (Figure 3F).

The two model-based agents, however, performed significantly better, with the influence zones algorithm outperforming DYNAQ (Figure 3F). Both learned to navigate to goal for more than half of the training trajectories and chose the shorter arm for >94% of these (Figure 3E,F). These results suggest that rapidly learning to navigate the arena environment with limited exploration requires a learning algorithm that goes beyond naïve model-free rules and incorporates elements such as internal replay or the topology of the environment.

## Discussion

This work shows that mice in a novel environment learn to choose shortest escape route to shelter when there is more than one option. This learning process is fast and happens during spontaneous exploration, before mice have experienced any threats. The choice is done by selecting the route with the shortest path length and shortest angle to shelter.

Minimizing the path length during escape agrees with the straight flight trajectories observed in open arenas and appears to serve the purpose of minimizing exposure to danger. While in an open arena the shortest path is the direct one, here mice had to learn the path lengths of the different routes to choose the shortest path. The minimization of shelter angle is in line with the observation that mice keep track of a vector to the shelter in open arenas [2,7]. In the arenas tested in this work, mice seem to keep track of this vector and use it as a variable for choosing escape paths even though there was never a direct route to shelter available. While this decision strategy does not minimize exposure to risk in these arenas, selecting the path with the smallest shelter angle minimizes the Euclidean distance to the shelter. Should a shortcut suddenly appear along the escape path, following a default policy of moving closer to the shelter might provide an advantage.

While it may seem trivial that animals would choose the fastest escape route, this need not be the case. Escape strategies in the animal kingdom are diverse and there is often an advantage to using alternative strategies, such as outpacing the predator while not revealing where shelter is [4,19]. Perhaps surprisingly, we found that escape path choice was probabilistic despite the relative path lengths being fixed for each experimental session. This could reflect imperfect learning of the environment geometry, noise in the sensory and decision-making systems, or the effect of unmeasured variables. Alternatively, it could also provide an advantage by maintaining some amount of exploration while exploiting the fastest known route, or by increasing the unpredictability of the escape trajectory, which has been suggested to be advantageous for many species [4,20–22].

Our results highlight the close interplay between learning over two distinct timescales to generate adaptive behavior: individual experience and evolution. In our experiments mice had to learn the spatial properties of the arena, which they did through their natural drive and behavior. Even though no task structure nor explicit instructions were provided, all animals explored the arenas efficiently, identifying the shelter, available paths, and extracting relevant arena features such as path length and angle to shelter. They then immediately relied on the estimation of path length and direct shelter vector to assume that minimizing these is the best value for escape. Thus, in addition to not having to learn that small, enclosed places offer shelter from threat, mice also did not have to learn the value of the different escape routes through experience of threat, or any form of punishment associated with taking the longer routes (e.g.: being exposed to an unpleasant stimulus for longer). Instead, they have an innate policy that guides escape path selection upon the first encounter with threat, thus removing the need for trial-and-error learning in a scenario in which errors could be fatal. This finding is in agreement with mice not needing to be exposed to threat to learn the direct shelter vector [2] or even a more complex subgoal shelter route [3] and suggests that mapping escape value onto the spatial environment is a priority of naturally behaving mice. A likely explanation is that, as prey species, during free exploration mice give high value to sheltering locations and routes that lead to these safe places, even in the absence of explicit threat. They extract relevant knowledge from the environment when it is safe to do so and identify a set of possible defensive actions. When threat does come and defensive actions need to be selected, an innate heuristic is leveraged to assign value to each alternative with no need for further learning. This combination of acquired and innate knowledge ensures that the animals can discover and select the most adaptive defensive action in the safest way, thus maximizing their chances of survival. From a neurobiology perspective, these findings provide a new paradigm for investigating the mechanisms of value assignment, how the estimation of spatial properties is linked to route decisions and how learned and innate information are integrated to guide decision-making.

A key finding in this study is that route learning was fast and required minimal exploration. This builds on previous work showing fast learning of shelter location [2,8,23] and supports recent findings showing that mice rapidly learn to navigate a maze for reward through natural exploration [11]. A picture that emerges from these studies is that natural exploration of space is the fundamental way that mice learn about the environment and express their behavioral choices, and therefore evolution has ensured that spatial learning is fast and prioritizes supporting survival needs. While it is unclear what learning algorithms mice use in the settings explored here, our reinforcement learning modeling suggests that a simple model free algorithm is not sufficient to generate the observed behavior. Instead, a more sophisticated learning process seems to be required to extract the necessary information from very limited experience. Alternatively, a simple learning algorithm could act on prior knowledge that is useful for solving the problem, such as a model for quickly estimating distances from self-motion. Our results invite future work investigating the biological basis of how innate and acquired knowledge interact to generate behavior, as well as on how abstractions of this interaction can be leveraged for developing efficient learning algorithms for machine learning applications [24].

## Experimental Procedures

### Resource availability

#### Lead contact

Further information and requests for resources and reagents should be directed to and will be fulfilled by the lead contact, Tiago Branco.

#### Materials availability

All arena design files are available upon request.

#### Data and code availability

All data reported in this paper will be shared by the lead contact upon request.

Analysis code can be found at the following GitHub repository: https://github.com/FedeClaudi/EscapePathSelection.

Any additional information required to reanalyze the data reported in this paper is available from the lead contact upon request.

### Experimental model and subject details

#### Animals

Male adult C57BL/6J Mice (8-12 weeks old) were single housed on a 12h light cycle and tested during the light phase. Mice from the same litter were randomly assigned to different experiments when possible. Most animals (78/119) were only used once, and the remaining were used for experiments on two arenas on different days. Mice that were used twice did not show any difference in escape response or arm choice probability compared to animals that were used only once. All experiments were performed under the UK Animals (Scientific Procedures) act of 1986 (PPL 70/7652 and PFE9BCE39).

### Methods details

#### Behavioral arena

The behavioral arenas consisted of white acrylic platforms elevated 30cm from the floor. Each arena was composed of octagonal platforms (24cm in diameter) and connecting bridges of various lengths (10cm wide). For the experiment shown in Figure 2E, some bridge sections were fitted with a computer-controlled servo motor which rotated a 20cm long bridge section by 90 degrees in the downward direction and created a gap that the mice could not traverse. The servo motors were controlled with custom Arduino code and activated manually.

**Table 1.** Arenas properties. Lengths and angles (relative to shelter direction) of left and right paths in arenas 1-5. The geodesic and angles ratios were defined as: 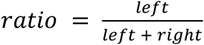.

|  | Arena 1 | Arena 2 | Arena 3 | Arena 4 | Arena 5 |
| --- | --- | --- | --- | --- | --- |
| **Left path length** | 505cm | 770cm | 872cm | 1085cm | 1085cm |
| **Right path length** | 505cm | 505cm | 505cm | 505cm | 1085cm |
| **Left path angle** | 45° | 45° | 45° | 45° | 45° |
| **Right path angle** | 45° | 45° | 45° | 45° | 90° |
| **Geodesic ratio** | 0.50 | 0.60 | 0.63 | 0.68 | 0.5 |
| **Angles ratio** | 0.50 | 0.50 | 0.50 | 0.50 | 0.33 |

#### Auditory and visual stimulation

Mice were presented with auditory stimuli consisting of three frequency modulated sweeps from 17 to 20 kHz of 3 seconds each at a sound pressure of 70-85dB as measured at the arena floor. In some experiments overhead visual stimuli were used. These were projected onto a screen positioned 1.8m above the arena floor and consisted of a dark circle (Weber contrast = −0.98) expanding over a period of 250ms [2]. The visual stimulus was repeated five times in short sequence with an interval of 500ms between repeats. No difference in behavior was observed between auditory and visual stimuli and therefore the data were pooled. Stimuli were triggered manually and controlled with software custom-written in LabVIEW (2015 64-bit, National Instrument). While manual stimulation could be a source of bias, the arena design ensured that mice would be in the same position and similar orientation across trials and experiments. No systematic difference in position or orientation was observed based on the selected escape path. The sound was played from the computer through an amplifier (TOPAZ AM10, Cambridge Audio) and speaker (L60, Pettersson). The audio signal was fed in parallel through a breakout board (BNC-2110, National Instruments) into a multifunction I/O board (PCIe-6353, National Instruments) and sampled at 10 KHz. To synchronize the audio and video, this signal was compared to the 30/40 Hz pulse triggering video frame acquisition, which was also fed as an input to the input/output board and sampled at 10 KHz. The visual stimuli and video were synchronized using a light dependent resistor whose voltage output depends on the amount of light it detects and thus reflected the presence/absence of visual stimuli. The resistor’s output was fed as input to the input-output board and sampled at 10 KHz.

#### Behavioral assay

Experimental sessions were filmed at 30 or 40fps with an overhead camera (positioned 1.8m above the arena floor). At the start of each experimental session mice were allowed to freely explore the arena for a period of ~10 minutes, during which they spontaneously found the shelter. After the exploration period, threats (either auditory or visual) were presented repeatedly while the animals were on a designated threat platform and facing in the direction opposite from the shelter platform. A stimulus response was considered an escape if the mouse reached the shelter within 10 seconds from stimulus onset. The number of trials in each experimental session varied across mice. Experiments were terminated when mice either remained in the shelter continuously for 30 minutes or failed to escape in response to three consecutive stimuli. Some experiments were performed in total darkness (with auditory stimuli only). No difference in behavior was observed between lights on and lights off experiments and thus the two datasets were pooled.

#### Animal tracking

The position and orientation of mice in the arena was reconstructed using DeepLabCut [25] to track the position of several body parts (snout, neck, body and tail base) in all recorded videos with a custom trained network. Post processing of tracking data included median filtering (window width: 7 frames), removal of low-confidence tracking (likelihood < 0.995) and affine transformation to a standard template arena to facilitate comparison across experiments [3]. Processed tracking data were stored in a custom DataJoint database [26] which also stored experimental metadata (e.g., mouse number, arena type, stimuli times etc.) and was used for all subsequent analysis.

#### Analysis code

All analysis was carried out using custom Python code and used several software packages from Python’s scientific software ecosystem: NumPy [27], Matplotlib [28], Scikit [29], Pandas [30], OpenCV [31], and StatsModels [32]. To calculate the animal’s orientation, we computed the vectors between the tail base and body and between the body and snout body part location as reconstructed by DeepLabCut, we then took the average of the two (we found this to be more stable than either vector alone). We set the shelter direction to be at 0 degrees in allocentric coordinates.

#### Reinforcement learning modelling

Three classes of Reinforcement Learning (RL; [17]) models were trained to navigate a grid world representation of the Arena 4 from the experimental study. All RL simulation, analysis and data visualization work were done in custom Python code. The grid world representation of Arena 4 consisted of a 50×50 array of quadrilateral cells with zeros corresponding to locations on the arena (126 cells in total) and ones to locations inaccessible by the agent. Agents could move in 8 cardinal directions (up, up-right, right, down-right, down, down-left, left, up-left) by one cell at the time and had to learn how to navigate the environment from a starting location to a goal location (corresponding to the threat and shelter locations in Arena 4 correspondingly). All agents were awarded a reward of 1 for reaching a cell < 3 cells distant from the goal location and received a penalty of −0.1 for attempting a move leading towards an inaccessible cell (upon which the agent did not move). For the QTable agent only (described below) a small (1e-8) reward was also delivered for any training step in which the agent moved to a new cell to encourage exploration.

#### Reinforcement learning models

Three different reinforcement learning model were used. All models shared the same environment (state space), actions space and reward function (with the exception of QTable as noted above). The three models were: QTable (model free RL; [17]), DYNA Q (model based; [17]) and Influence Zones (IZ, model based; [18]). All three models were implemented in custom python code. For all simulations the following parameters were used:

**Table 2.** Reinforcement learning agents parameters. Parameter values for the reinforcement learning agents.

|  | DISCOUNT | LEARNING RATE | EXPLORATION<br>RATE | EXPLORATION<br>RATE DECAY |
| --- | --- | --- | --- | --- |
| **QTABLE** | 0.8 | 0.9 | 0.4 | 0.995 |
| **DYNA Q** | 0.8 | 0.8 | 0.4 | 0.995 |
| **INFLUENCE ZONES** | 0.8 | 0.2 | 0.4 | 0.995 |

The DYNAQ model includes a planning step in which randomly sampled entries from the agent’s model are used to update the value function. In all simulations the number of samples used for each planning step was set to 20.

The IZ model had additional parameters. These include a TD error threshold for one-step updates of value function (1e-10) and a threshold for n-step updates (0.1) and additional parameters for the Instantaneous Topological Map (ITM; [33]) model used by IZ: ITM learning rate (0.2) and max error (1.2).

#### Free exploration training

In the free exploration training regime, agents were trained for 250 episodes of maximum 500 training steps each. For each training episode the agent was initialized in a random location on the grid world arena and had to navigate to the goal location. The episode terminated when the agent took 500 steps or if it reached the goal location. During training at each step agents selected the action to perform using an epsilon greedy strategy: a random action is chosen with probability equal to the exploration rate parameter, otherwise the action with the current highest value is selected. At the end of each episode, the exploration rate decayed by a factor set by the parameter exploration rate decay.

To assess the agent’s performance during learning, at the end of each training episode the agent was initialized at the start location and allowed to act greedily (i.e., with no randomly selected actions). If the agent reached the shelter location the simulation was marked successful, otherwise it was labelled as a failure. If the agent attempted an illegal move (i.e., trying to move to an inaccessible cell) the simulation was terminated and considered a failure. The agents were not allowed to use the experience from this evaluation simulation for learning.

#### Guided exploration training

In the guided exploration training, RL agents followed the exploration trajectory from the experimental animals. Tracking data from each experiment’s exploration phase was registered to the grid world arena through an affine transformation (scaling and shift). Tracking data was represented at a higher spatial resolution that the grid world arena and did not match the grid world arena layout perfectly (due to imperfect registration of the tracking data to the standard template). The first issue was resolved by assigning, for each frame in the tracking data, the grid world arena cell closest to it. Imperfect alignment and tracking errors could not be corrected in some experimental sessions and these were discarded, leaving 42 valid sessions. As mice often remained in the same location for extended periods of time during natural exploration (e.g., in the shelter), these periods were eliminated from the tracking data and only frames in which the mouse moved from one arena cell to another were kept. The tracking data was then used to guide the movement of all agents during the training phase. For a given session’s data, the agent was initialized at the arena location corresponding to the first frame in the tracking data. For each step, the location of the next arena cell was identified and the action leading from the agent’s current cell to the next was identified. The agent then performed the selected action, experienced rewards and learned, as it would have during free exploration. Thus, the main difference between the free and guided exploration paradigms was that in the guided exploration regime agents were not allowed to select which action to perform as this was determined by the tracking data.

Once the agent followed each step from the tracking data the training phase was concluded. The agent was then initialized at the start location and allowed to act greedily following the value function it learned during training, with the goal of reaching the shelter location. If the goal was reached the simulation was classified as successful, otherwise it was classified as a failure. If the agent attempted an illegal move (i.e., trying to move to an inaccessible cell) the simulation was terminated and classified as a failure.

### Quantification and statistical analysis

#### Quantification of escape probability

To calculate the probability of escaping in response to the threat stimulus, the movement trajectories during the first 10s after stimulus onset were analyzed, and the fraction of trajectories terminating on the shelter platform was computed. For comparison, randomly selected time points in which the same animals where on the threat platform but not presented with a threatening stimulus were selected, and the fraction of shelter arrivals in this random sample was estimated too. The number of randomly selected time points matched the number of trials. Fisher’s exact test was used to determine the significance of the difference in number of shelter-arrival trajectories between stimuli and control groups.

#### Quantification of the probability of escaping along a dead-end

To determine the probability of escape along the dead-end arm in arena 1, all trials from experiments in the arena were pooled. The trajectory in the 10 seconds following threat presentation was analyzed to determine which arm the mouse first moved to after leaving the threat platform, and the probability of escape along each of the three arms (left, right and dead end) was then computed. To distinguish between trials in which the mice escaped vs trials in which the mouse ignored the stimulus but still moved away from the threat platform, we used two criteria. First, since escapes are characterized by a higher running speed than normal locomotion [2], we only included in the analysis trials in which the mouse was moving at a speed higher than 35 cm/s when it left the threat platform. Second, as escapes have fast reaction times and mice leave the threat platform within the first 3 seconds from stimulus, only trials in which the mouse left the threat platform within 4 seconds from stimulus onset were considered escapes.

#### Quantification of heading direction

To quantify the average heading direction during escape, movement trajectories from each arena were selected and grouped into left or right path escapes. The trajectories where then truncated to the frame in which the mouse left the threat platform and their duration normalized. The average heading direction at regular intervals across all trajectories in the same group was then computed.

#### Quantification of arm choice probability

To estimate the probability of escape along a given path, all trials from experiments on each arena were pooled and the probability of selecting the right or left arm was computed. In experiments in which the arena design changed during the course of an experiment, trials were pooled across animals and grouped into baseline (before the change) and ‘flipped’ (after the change). Fisher’s exact test was used to determine whether the number of escapes along the right path differed between the baseline and flipped conditions.

In addition to estimating the probability of selecting an arm, the posterior distribution of the probability value was estimated using a Bayesian model. The model had a Beta distribution prior (parameters: a=1, b=1) and a Binomial distribution as likelihood (n = total number of trials, k = total number of right path escapes). The resulting posterior distribution is then a Beta distribution whose parameters are given by:

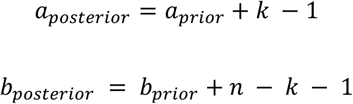

To compare the probability of selecting the right arm between naïve and experienced trials, for each arena the first escape trial for each mouse was classified as naïve while all other trials were classified as experienced. When a mouse was used for more than one experiment, only the first trial on the first experiment the mouse was used for was considered naïve. The probability of escaping along the right path was computed for the naïve trials. Because the number of experienced trials is larger than the naïve condition, to compare the probability of taking the right path between the two groups, experienced trials were randomly sampled without replacement to match the number of naïve trials in the same arena, and the probability of taking the right path was then computed. This procedure was repeated 10 times to generate a distribution of probability values, and the mean and 95^th^ percentile interval were computed from this distribution.

#### Arm choice probability for naïve vs experienced trials

To determine whether mice required repeated experience with threat to select the preferred escape path we identified the very first stimulus presentation of each animal. We grouped such “naïve” trials across individuals tested on the same experimental arena and estimated the probability of escape along the right path in this subset of the data. To compare naïve vs experienced (i.e. following the first encounter with threat) trials, we randomly sampled from the experienced trials from each arena matching the number of naïve trials in the same arena and we computed the probability of escape on the right arm. We repeated the sampling procedure 100 times to obtain a distribution of probability values for different random subsets.

#### Change in path preference with time

To quantify the change in probability of escape along the right path over time, for each arena we pooled all trials that occurred during the first 60 minutes from the start of the experiment. We then binned the trials based on the time since experiment start (interval between bins: 120 seconds, bin width: 300s) and computed the posterior distribution of p(R) as described above.

#### Quantification of shelter-threat trips during exploration

To quantify the number of trips between the shelter and the threat platform during exploration, the tracking trajectory corresponding to the exploration period (from start of the experiment to one frame before the first stimulus) was analyzed. For each frame, the mouse was assigned to one of four regions of interest (‘shelter’, ‘right arm’, ‘threat’, ‘left arm’) based on the coordinates of the tracking data registered to a standard template image as described above. A trip was ended when the mouse arrived at the shelter (threat) platform and started at the last frame in which the mouse was on the threat (shelter) platform. Incomplete trips (e.g.: the mouse left the shelter platform and returned to it without first reaching the threat platform) were discarded.

#### Predicting escape arm with GLM

To predict the probability of escaping along the right arm from trial data, a binomial generalized linear model (GLM) with a logit link faction was used (implemented in StatsModels; [32]). All trials from all arenas were pooled and split between train and test sets (stratified k-fold repeated cross-validation; five different splits of the trials were used, and the data were split such that the test set was roughly balanced with respect to the number of trials from the corresponding arena; this procedure was repeated four times with different random splits each time, yielding a total of 20 model fits). The GLM model attempted to estimate the probability of escape along the right path for each trial based on: 1) the geodesic ratio of the trial’s arena, 2) the angles ratio of the trial’s arena, 3) the trial time (in seconds) since the start of the experimental session, 4) the identity of the origin arm. Categorical variables were one-hot encoded, and all variables were normalized to the 0-1 range. The accuracy of the model’s predictions on the test data was estimated with the Pearson’s correlation between the predicted probability of escape on the right arm and the arm chosen by the animal. This accuracy measure was compared to the accuracy of models fitted on randomly shuffled data. The full model described above was then compared to models lacking one or two of the input variables to estimate the effect of each variable. Each model was fitted on k-fold cross validated data (k=5) and the procedure was repeated four times using different random number generator seeds for each repeat. The coefficient weights of each parameter for each fit of the full model were used to estimate the average and standard deviation of the coefficient weights.

#### Decoding escape path from threat trajectory

To decode the escape arm from the trajectory on the threat platform we used a logistic regression model (implemented in Scikit; [29]). Trajectories from all trials in each experimental arena were pooled and their duration was normalized. To assess how the model performed as mice moved away from the threat location and towards the escape arms, 8-9 time points were selected corresponding to different average positions along the axis between the threat and shelter platform. For each time point, the animal’s average orientation in the five frames after the time point and the trial’s escape arm were used. The data were randomly split between a training and test set (test set 0.33% of the trials) and the training set was used to fit the model to predict the escape arm based on the orientation value. The model’s accuracy score on held out training data was then computed. The procedure was repeated 100 times for each time point with a different random split of the data and the average accuracy computed.

#### Quantification of RL models performance

To quantify the performance of RL agents trained to navigate the grid world arena under the free exploration regime, we trained 64 repetitions of each model. At the end of each training episode, each repetition was tested on its ability to navigate to the goal location and returned a 1 for successes and 0 otherwise. Thus, a vector of outcomes was constructed based on the value returned by each repetition and the overall score was given by the mean and standard error of the mean (SEM) of the outcomes vector. For visualization, the mean and SEM accuracy at each training episodes were displayed following smoothing with a rolling mean filter with window width of 6 episodes.

To quantify the performance of RL agents trained under the guided exploration regime, we trained 10 repetitions of each RL model on the tracking data from each experimental session. Under this regime, unlike in free exploration, training is fully deterministic because the actions are specified by the tracking data, the only variability emerges from the DYNA-Q model’s probabilistic sampling of its model at each training step. After training, all repetitions of each model were tested on their ability to navigate from the start to the threat location. The model was considered successful if at least 8/10 repetitions successfully reached the goal location.

To produce the state occupancy heatmaps in Figure 3B we recorded all cell visits for one example agent trained under the free exploration regime and one example agent trained under the guided exploration regime. The total number of visits to visits to each cell was then used to produce the heatmap. To visualize the preferred action at each cell (Figure 3E) we trained one example agent for each class of RL algorithm and training regime and displayed the action with highest value for each cell.

## Author Contributions

F.C. and T.B. designed the study and experiments and wrote the manuscript. F.C. performed all experiments and analyses.

D.C. contributed to experiment design, data analysis design and interpretation.

## Acknowledgments

This work was funded by a Wellcome Senior Research Fellowship (214352/Z/18/Z) and by the Sainsbury Wellcome Centre Core Grant from the Gatsby Charitable Foundation and Wellcome (090843/F/09/Z) (T.B.), and the SWC PhD Programme (F.C.). We thank members of the Branco lab for discussions, and Ruben Vale and Phillip Shamash for comments on the manuscript.

## Supplementary Figures

**Figure S1.**
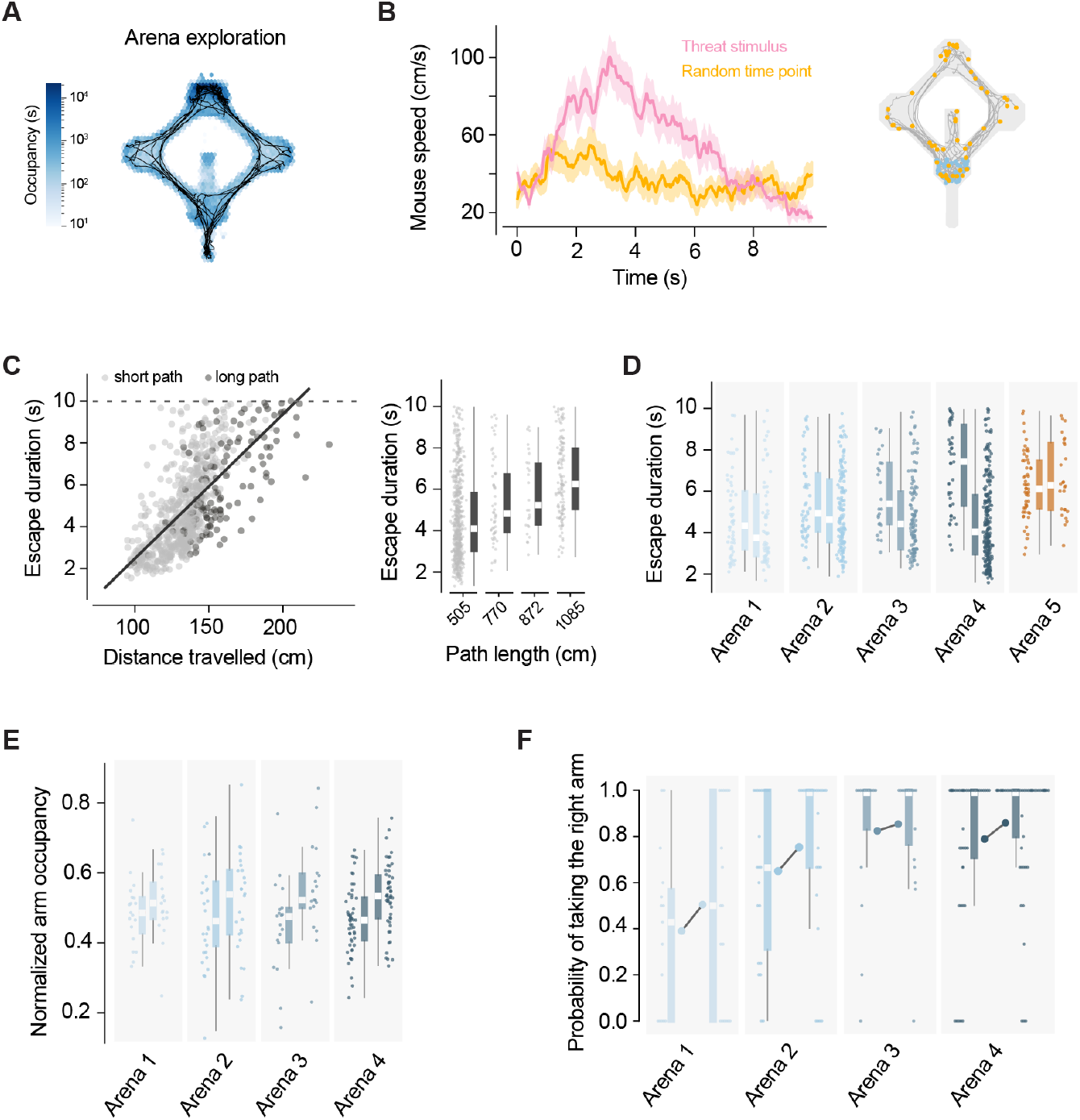
Escape duration correlates with distance and route choice does not depend on path exploration. **(A)** Heatmap of arena occupancy during exploration of Arena 1 with example tracking traces from one mouse (black). **(B)** Left, speed trajectory in the 10s following the threat stimulus (pink) or a randomly selected time point when the mouse was in the threat area (yellow). Probability of reaching the shelter after threat stimulus = 0.65 vs 0.17 for random time points (p = 0.00097, Fisher exact test). Bold curves and shaded areas show mean and standard deviation across trials. Right, tracking traces for the random time point condition, blue circles show the starting location, orange circles the final location. **(C)** Left, correlation between distance travelled during escape and escape duration. Each dot represents a single escape trial, darker dots are trials in which the animals selected the longer paths (in Arenas 2,3,4). Black line shows best fit of linear regression (slope: 0.07, intercept: −4.4). Right, escape duration along paths of different lengths. Each dot represents a single trial, box plots show the median and 95th percentile of the distribution. **(D)** Escape duration for left and right path escapes across arenas (for each arena the left bar shows escapes on the left and the right bar shows escapes on the right). Each dot represents a single trial, box plots show the median and 95th percentile of the distribution (t-test with Bonferroni correction for comparison between left and right: Arena 1, p=1.0; Arena 2, p=0.5; Arena 3, p=0.04; Arena 4, p=1.9e-12; Arena 5, p=1.0). **(E)** Fraction of time spent on left and right path during exploration normalized by the relative length of each path (shown by bars on the left and right for each arena, respectively). Circles indicate individual mice, box plots show the median and 95th percentile of the distribution (t-test with Bonferroni correction for comparison between left and right: Arena 1, p=1.0; Arena 2, p=1.0; Arena 3, p=1.0; Arena 4, p=0.002; Arena 5, p=1.0). **(F)** Probability of escape on the right path for subsets of trials in which mice reached the threat platform from the left or the right path (shown by bars on the left and right for each arena, respectively). Colored circles represent individual mice, box plots show the median and 95th percentile of the distribution (values between left are not significantly different for all arenas).

**Figure S2.**
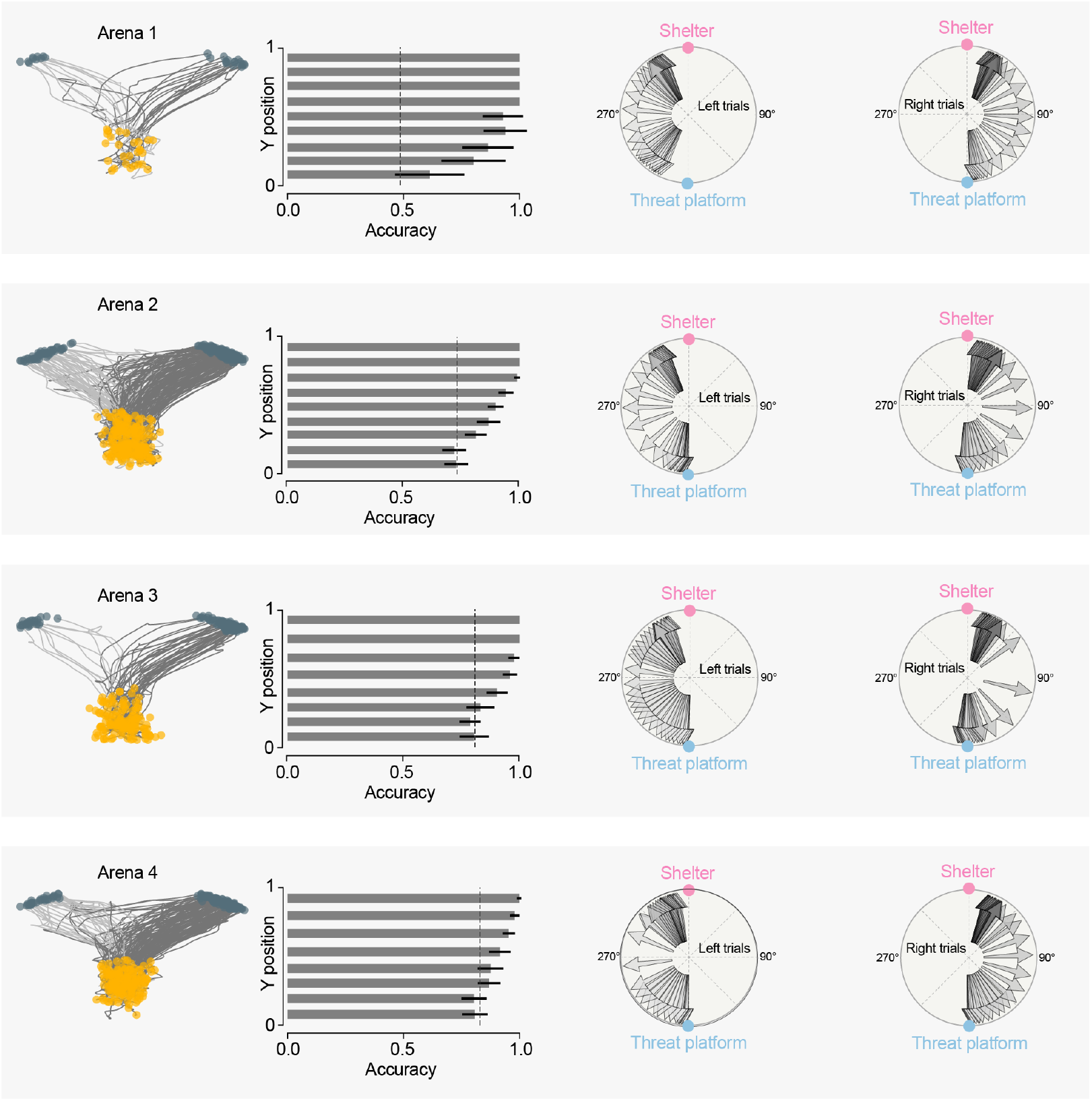
Escape path choice can be decoded from the movement trajectory in the threat platform. Decoding of escape path selection probability based on tracking data on the threat platform. Left, tracking traces for all trials in each arena. Darker traces show escapes on the right path, orange circles the initial position and blue circles the animal’s position at the time of leaving the threat platform. Center, test set accuracy of a logistic model predicting escape path choice based on tracking data binned along the threat-shelter axis. Right, average orientation across trials during escape progression for left and right trials. Arrows show orientation for each time bin, darker colors indicated later time points.

## References

1. Evans, D.A., Stempel, A.V., Vale, R., Ruehle, S., Lefler, Y., and Branco, T. (2018). A synaptic threshold mechanism for computing escape decisions. Nature 558, 590–594.

2. Vale, R., Evans, D.A., and Branco, T. (2017). Rapid Spatial Learning Controls Instinctive Defensive Behavior in Mice. Curr. Biol. 27, 1342–1349.

3. Shamash, P., Olesen, S.F., Iordanidou, P., Campagner, D., Banerjee, N., and Branco, T. (2021). Mice learn multistep routes by memorizing subgoal locations. Nat. Neurosci. Available at: http://dx.doi.org/10.1038/s41593-021-00884-8.

4. Cooper, W., Jr, Cooper, W.E., Jr, and Blumstein, D.T. (2015). Escaping From Predators: An Integrative View of Escape Decisions (Cambridge University Press).

5. De Camp, J.E. (1920). Relative distance as a factor in the white rat’s selection of a path. Psychobiology 2, 245.

6. Snygg, D. (1935). Mazes in Which Rats Take the Longer Path to Food. J. Psychol. 1, 153–166.

7. Vale, R., Campagner, D., Iordanidou, P., Arocas, O.P., Tan, Y.L., Vanessa Stempel, A., Keshavarzi, S., Petersen, R.S., Margrie, T.W., and Branco, T. (2020). A cortico-collicular circuit for accurate orientation to shelter during escape. bioRxiv, 2020.05.26.117598. Available at: https://www.biorxiv.org/content/10.1101/2020.05.26.117598v1 [Accessed June 2, 2020].

8. Yilmaz, M., and Meister, M. (2013). Rapid innate defensive responses of mice to looming visual stimuli. Curr. Biol. 23, 2011–2015.

9. Mongeau, R., Miller, G.A., Chiang, E., and Anderson, D.J. (2003). Neural correlates of competing fear behaviors evoked by an innately aversive stimulus. J. Neurosci. 23, 3855–3868.

10. Ellard, C.G., and Eller, M.C. (2009). Spatial cognition in the gerbil: computing optimal escape routes from visual threats. Anim. Cogn. 12, 333–345.

11. Rosenberg, M., Zhang, T., Perona, P., and Meister, M. (2021). Mice in a labyrinth show rapid learning, sudden insight, and efficient exploration. Elife 10. Available at: http://dx.doi.org/10.7554/eLife.66175.

12. Eason, P.K., Nason, L.D., and Alexander, J.E., Jr. (2019). Squirrels Do the Math: Flight Trajectories in Eastern Gray Squirrels (Sciurus carolinensis). Available at: https://www.frontiersin.org/article/10.3389/fevo.2019.00066/full [Accessed June 4, 2020].

13. Tolman, E.C. (1948). Cognitive maps in rats and men. Psychol. Rev. 55, 189–208.

14. International Brain Laboratory, Aguillon-Rodriguez, V., Angelaki, D., Bayer, H., Bonacchi, N., Carandini, M., Cazettes, F., Chapuis, G., Churchland, A.K., Dan, Y., et al. (2021). Standardized and reproducible measurement of decision-making in mice. Elife 10. Available at: https://elifesciences.org/articles/63711.

15. Arulkumaran, K., Deisenroth, M.P., Brundage, M., and Bharath, A.A. (2017). A Brief Survey of Deep Reinforcement Learning. arXiv [cs.LG]. Available at: http://arxiv.org/abs/1708.05866.

16. Silver, D., Hubert, T., Schrittwieser, J., Antonoglou, I., Lai, M., Guez, A., Lanctot, M., Sifre, L., Kumaran, D., Graepel, T., et al. (2018). A general reinforcement learning algorithm that masters chess, shogi, and Go through self-play. Science 362, 1140–1144.

17. Richard R. Sutton, A.G.B. (2015). Reinforcement Learning - An introduction. The MIT Press. Available at: https://web.stanford.edu/class/psych209/Readings/SuttonBartoIPRLBook2ndEd.pdf.

18. Braga, A.P. de S., and Araújo, A.F.R. (2006). Influence zones: A strategy to enhance reinforcement learning. Neurocomputing 70, 21–34.

19. Mattingly, W.B., and Jayne, B.C. (2005). The choice of arboreal escape paths and its consequences for the locomotor behaviour of four species of Anolis lizards. Anim. Behav. 70, 1239–1250.

20. Domenici, P., Booth, D., Blagburn, J.M., and Bacon, J.P. (2008). Cockroaches keep predators guessing by using preferred escape trajectories. Curr. Biol. 18, 1792–1796.

21. Domenici, P., Blagburn, J.M., and Bacon, J.P. (2011). Animal escapology I: theoretical issues and emerging trends in escape trajectories. J. Exp. Biol. 214, 2463–2473.

22. Domenici, P., Blagburn, J.M., and Bacon, J.P. (2011). Animal escapology II: escape trajectory case studies. J. Exp. Biol. 214, 2474–2494.

23. De Franceschi, G., Vivattanasarn, T., Saleem, A.B., and Solomon, S.G. (2016). Vision Guides Selection of Freeze or Flight Defense Strategies in Mice. Curr. Biol. 26, 2150–2154.

24. Zador, A.M. (2019). A critique of pure learning and what artificial neural networks can learn from animal brains. Nat. Commun. 10, 3770.

25. Mathis, A., Mamidanna, P., Cury, K.M., Abe, T., Murthy, V.N., Mathis, M.W., and Bethge, M. (2018). DeepLabCut: markerless pose estimation of user-defined body parts with deep learning. Nat. Neurosci. 21, 1281–1289.

26. Yatsenko, D., Reimer, J., Ecker, A.S., Walker, E.Y., Sinz, F., Berens, P., Hoenselaar, A., James Cotton, R., Siapas, A.S., and Tolias, A.S. (2015). DataJoint: managing big scientific data using MATLAB or Python. bioRxiv, 031658. Available at: https://www.biorxiv.org/content/10.1101/031658v1.abstract [Accessed July 28, 2021].

27. Harris, C.R., Millman, K.J., van der Walt, S.J., Gommers, R., Virtanen, P., Cournapeau, D., Wieser, E., Taylor, J., Berg, S., Smith, N.J., et al. (2020). Array programming with NumPy. Nature 585, 357–362.

28. Hunter, J.D. (2007). Matplotlib: A 2D Graphics Environment. Computing in Science Engineering 9, 90–95.

29. Pedregosa, F., Varoquaux, G., Gramfort, A., Michel, V., Thirion, B., Grisel, O., Blondel, M., Prettenhofer, P., Weiss, R., Dubourg, V., et al. (2011). Scikit-learn: Machine Learning in Python. J. Mach. Learn. Res. 12, 2825–2830.

30. Reback, J., McKinney, W., jbrockmendel, Van den Bossche, J., Augspurger, T., Cloud, P., gfyoung, Sinhrks, Klein, A., Roeschke, M., et al. (2020). pandas-dev/pandas: Pandas 1.0.3 Available at: https://zenodo.org/record/3715232.

31. Bradski, G., and Kaehler, A. (2000). OpenCV. Dr. Dobb’s journal of software tools 3. Available at: http://roswiki.autolabor.com.cn/attachments/Events(2f)ICRA2010Tutorial/ICRA_2010_OpenCV_Tutorial.pdf.

32. Seabold, S., and Perktold, J. (2010). Statsmodels: Econometric and statistical modeling with python. In Proceedings of the 9th Python in Science Conference (SciPy). Available at: http://conference.scipy.org/proceedings/scipy2010/pdfs/seabold.pdf [Accessed July 29, 2021].

33. Jockusch, J., and Ritter, H. (1999). An instantaneous topological mapping model for correlated stimuli. In IJCNN’99. International Joint Conference on Neural Networks. Proceedings (Cat. No.99CH36339), pp. 529–534 vol.1.

